# Morpholino-mediated knockdown of the brain mineralocorticoid receptor affects glucocorticoid signaling and neuroplasticity in wild ocellated wrasse (*Symphodus ocellatus*)

**DOI:** 10.1101/2021.11.17.468986

**Authors:** Bridget M. Nugent, Kelly A. Stiver, Jiawei Han, Holly K. Kindsvater, Susan E. Marsh-Rollo, Hans A. Hofmann, Suzanne H. Alonzo

## Abstract

Uncovering the genetic, physiological, and developmental mechanisms underlying phenotypic variation is necessary for understanding how genetic and genomic variation shape phenotypic variation and for discovering possible targets of selection. Although the neural and endocrine mechanisms underlying social behavior are evolutionarily ancient, we lack an understanding of the proximate causes and evolutionary consequences of variation in these mechanisms. Here, we examine in the natural environment the behavioral, neuromolecular, and fitness consequences of a morpholino-mediated knockdown of the mineralocorticoid receptor (MR) in the brain of nesting males of the ocellated wrasse, *Symphodus ocellatus*, a species with male alternative reproductive tactics. Even though MR knockdown did not significantly change male behavior directly, this experimental manipulation strongly altered glucocorticoid signaling and neuroplasticity in the preoptic area, the putative hippocampus homolog, and the putative basolateral amygdala homolog. We also found that individual variation in stress axis gene expression and neuroplasticity is strongly associated with variation in male behavior and fitness-related traits. The brain region-specific effects of MR knockdown on phenotypic integration in the wild reported here suggest specific neuroendocrine and neuroplasticity pathways that may be targets of selection.

## INTRODUCTION

The neural and endocrine mechanisms underlying social behavior are broadly conserved across vertebrates (Toth & Robinson 2007; O’Connell & Hofmann 2012; Weitekamp & Hofmann 2017), in part because the biological functions and metabolic needs that drive these behaviors are often similar (O’Connell & Hofmann 2011a; Hofmann et al. 2014). Despite using the same basic underlying neuroendocrine “building blocks,” the specific behavioral patterns produced by them can vary widely among and within species. The neural and molecular mechanisms that underpin this variation, and their consequences for evolutionary fitness, are not yet well understood. Yet, this is precisely the information needed to know how these mechanisms evolve and the ways in which they may bias, constrain, and even enable the diversity of social interactions we see in nature.

Because uncovering the genetic, physiological, and developmental mechanisms underlying phenotypic variation can be challenging, behavioral ecologists initially posited that such mechanisms do not constrain the evolutionary trajectory of a trait and thus do not need to be incorporated explicitly when studying its evolution and adaptive value (Lloyd, 1977; Grafen, 1984). This simplifying assumption has facilitated tremendous progress in our understanding of how selective pressures optimize a given phenotype to its environment. However, knowing the genes as well as the physiological and neural intermediates associated with a (behavioral) trait of interest is important for understanding how genetic and genomic variation shape phenotypic variation and how behavioral flexibility, phenotypic plasticity, and learning impinge on an animal’s fitness (Hofmann et al., 2014; Rittschof & Robinson, 2014). In addition, the study of the proximate mechanisms underlying behavior enables genotype-to-phenotype mapping and can provide insight into past selection and the evolutionary history of behavior. Finally, knowing the genetic and physiological underpinnings of behavior allows us to manipulate these phenotypes, which can uncover targets of selection. Here, we build this essential knowledge by examining the neuroendocrine mechanisms underlying social behavior in a marine fish, using a morpholino-mediated knockdown of the mineralocorticoid receptor (MR) in the brain and observing the behavioral, neuromolecular, and fitness consequences of this experimental manipulation of fish observed under natural conditions in the wild.

The vertebrate stress axis, or hypothalamic-pituitary-adrenal (interrenal in anamniotes; HPA/I) axis, is a highly conserved system that integrates relevant internal and external cues in response to an environmental stressor (i.e., any external condition that disrupts, or threatens to disrupt, homeostasis). Activation of this system in form of a “stress response” allows an individual to react to unpredictable environmental perturbations, for example by adjusting their energy expenditure and behavior in ways that are adaptive (Raulo & Dantzer 2018). The stress response mediated by the HPA/I axis is initiated by the release of corticotropin-releasing factor (CRF) from the hypothalamus, which signals to the pituitary to release adrenocorticotropic hormone (ACTH), which in turn signals the adrenal glands to release glucocorticoid steroid hormones (e.g., cortisol, CORT, in fishes) (Denver, 2009; Lowry and Moore, 2006; Wendelaar Bonga, 1997). Increases in CORT in response to a stressor act throughout the central nervous system and periphery on glucocorticoid (GR) and mineralocorticoid receptors (MR) to establish HPA homeostasis (Takahashi & Sakamoto, 2013). CORT binding to high affinity MRs in response to a stressor is thought to initiate the stress response, while GR signaling appears to be responsible for its adaptability (Joëls et al., 2008). In mammals, HPA dysregulation is commonly associated with alterations in social and sexual behaviors (Blanchard et al., 2001), as relatively small changes in HPA signaling can have profound effects on hormonal homeostasis and behavior. Although MR is widely accepted as an integral part of the mammalian stress axis, GR signaling appears to be more relevant to stress reactivity and behavior in mammals (Joëls & de Kloet, 2017). Conversely, MR – which is thought to be ancestral to GR (Bridgham et al., 2006) – appears to be the principal receptor for glucocorticoid signaling in teleosts (Greenwood et al., 2003; Stolte et al., 2006), yet to this day MR is one of the least studied hormone receptors, despite its likely implications for behavior. Interestingly, several studies have reported very high levels of MR expression in the brain of several teleost species (Greenwood et al., 2003; Sturm et al., 2005; Arterbery et al., 2010; Takahashi & Sakamoto, 2013). In addition, variation in HPA/I axis signaling is often considered to be adaptive in a species-specific manner. For example, in some species, relatively high levels of CORT are associated with decreases in reproductive behavior, whereas in others CORT can facilitate such behavior (Raulo & Dantzer, 2018).

In addition to directing stress responses, neuroendocrine signaling mediates individual variation in social behavior across vertebrates (Adkins-Regan 2005; Goodson 2005; Rodgers et al. 2013, Raulo & Dantzer 2018), including in species with alternative reproductive tactics where differences in hormone signaling contribute to discrete differences in reproductive tactics and success (Knapp 2003; Nugent et al. 2016). Though sometimes the result of a genetic polymorphism, alternative reproductive tactics are often the consequence of phenotypic plasticity, where animals develop distinct behavioral and physiological attributes depending on the ecological and social conditions they encounter (Taborsky et al., 2008). At the level of the nervous system, these organismal changes are thought to be accompanied by the modification of synaptic connections between neurons (synaptic plasticity) and the birth of new neurons (neurogenesis), even in adult animals (Toda & Gage, 2018). Glucocorticoid signaling is linked to neural plasticity (McEwen, 2017) and CORT signaling may thus impact behavioral plasticity, perhaps as a mechanism allowing for rapid adaptation to social and environmental perturbations (Raulo & Dantzer, 2018). However, whether glucocorticoids act via MR or GR to alter neuroplasticity in the teleost brain is not known. The activity of genes involved in neural plasticity (i.e., changes in synaptogenesis and neurogenesis) can be used to assess this neuroplasticity. Prominent examples include doublecortin (DXC) as a marker for neurogenesis (Balthazart & Ball, 2016) and neurabin (also known as spinophilin in reference to its role in synaptic transmission and plasticity within the neuronal post-synaptic density) as a marker for synaptogenesis (Sarrouilhe et al., 2006). Similarly, brain-derived neurotrophic factor (BDNF) regulates neural plasticity (Gray et al., 2013), learning and memory (Ninan, 2014), and social cognition (Teles et al., 2016). While alternative reproductive tactics have been studied extensively, relatively little is known about the neural mechanisms and neuroplasticity underlying the striking differences in behavior these alternative phenotypes often exhibit.

Numerous brain regions have been shown to respond to glucocorticoid signaling as well as display neural plasticity in response to changes in the environment, especially also in the context of social behavior. The Social Decision-Making Network (SDMN) is a highly conserved fore- and midbrain circuit that is critical for evaluating stimulus salience and regulating sexual, aggressive and parental behavior across vertebrates (O’Connell & Hofmann 2011b; O’Connell & Hofmann 2012). It has been suggested that the diversity of vertebrate social behavior can be explained, in part, by variations on conserved spatial gene expression networks in the SDMN (O’Connell & Hofmann 2012). In addition, several SDMN nodes have been shown to play a critical role in HPA/I axis function. First, the **preoptic area (POA)** plays a fundamental role in regulating reproduction and social behavior across vertebrates (Dominguez & Hull 2005; Riters et al. 2004; Greenwood et al. 2008; Hattori & Wilczynski 2009; O’Connell et al. 2013; So et al. 2015). The paraventricular nucleus (PVN) within the POA contains numerous peptidergic cell populations, including those that encode CRF (which regulates the HPA/I axis: Denver, 2009; Lowry & Moore, 2006; Wendelaar Bonga, 1997), BDNF (which is upregulated in response to a stressor: Smith et al., 1995; and can increase CRF expression: Toriya et al., 2010), and arginine vasopressin (AVP, which regulates numerous aspects of social behavior: Oldfield et al, 2014; and can also co-activate the HPA/I axis: Aguilera & Rabadan-Diehl, 2000; Gesto et al., 2014). Second, the **hippocampus (HIP)** is critical for the formation of episodic memories and relational memory representations of the environment and/or experiences (O’Keefe & Nadel 1978; Andersen et al. 2007; Humphries & Prescott 2010), and numerous studies have demonstrated the importance of glucocorticoid signaling in this region (reviewed in Kim et al., 2015). The putative teleost homolog of the mammalian HIP, the pallial area Dl (O’Connell & Hofmann, 2011b) performs similar functions (Rodriguez et al., 2002; Elliott et al., 2017). Finally, the **basolateral amygdala (blAMY)** regulates affective and goal-directed behavior (Maeda and Mogenson 1981; LeDoux 2000; Moreno and González 2007) and is involved in social recognition (Wang et al., 2014) and social dominance (Dulka et al., 2018). Importantly, glucocorticoids activate blAMY neurons, thereby facilitating the consolidation of emotional memories (Duvarci & Paré, 2007). The putative teleost homolog of the mammalian blAMY, the pallial area Dm (O’Connell & Hofmann, 2011b), performs similar functions (Portavella et al., 2002). Together, examining neural activity and gene expression in POA, HIP, and blAMY – ideally in the natural environment – provides an excellent opportunity for understanding how the stress axis contributes to behavioral variation. Such an integrative approach demands a model system that shows considerable phenotypic variation (e.g., alternative reproductive tactics) and allows for detailed observation and experimental manipulation in its natural habitat.

The ocellated wrasse (*Symphodus ocellatus*) is a powerful system for conducting studies in the natural environment that connect variation in physiology to variation in social behavior and reproductive success (Dean et al., 2017; Nugent, Stiver, Alonzo, & Hofmann, 2016; Stiver, Harris, Townsend, Hofmann, & Alonzo, 2014; Nugent et al. 2016 and 2019). Three discrete male alternative types exist in this species (Alonzo, Taborsky, & Wirtz, 2000; Taborsky, Hudde, & Wirtz, 1987; Warner & Lejeune, 1985): Large, colorful **nesting males** are socially dominant, build and defend nests, court females, engage in paternal care, and have high levels of circulating androgens. Small, parasitically breeding **sneaker males** do not engage in courtship or paternal care, and instead opportunistically spawn in the nests of nesting males (Alonzo, 2004; Taborsky et al., 1987; Warner & Lejeune, 1985). **Satellite males** form an intermediate type whose presence is tolerated by the nesting male, as satellites help by chasing away sneaker males and attempt to bring females to the nest, but they do not engage in parental care (Stiver & Alonzo, 2013; Taborsky et al., 1987). This tolerance by the nesting male allows satellite males to sneak spawn when the nesting male is distracted (Stiver & Alonzo, 2013). During their breeding season, nesting males guard, tend and spawn in their nests and are highly aggressive towards conspecific males and heterospecific egg predators (Lejeune, 1985). In the ocellated wrasse, sperm competition plays a critical role in determining male reproductive success. Sneak spawning is rampant at active nests, with cuckoldry occurring at 100% of sampled nests (Alonzo & Heckman, 2010). Sneaker males pose a large threat to the reproductive success of nesting males since they release more sperm per spawn compared to the other male morphs, and the number of sneakers at a nest directly correlates with the intensity of sperm competition (Alonzo & Warner, 2000). However, since sneakers and satellite males do not engage in paternal care, their reproductive success relies on parental care by the nesting male. The behavior of every single individual in the group can therefore have marked fitness consequences for the entire social group breeding at a nesting site, but the behavior of the nesting male is of particular importance (Stiver et al., 2019).

We have previously shown that variation in the HPI axis is associated with the behavioral and physiological differences between *S. ocellatus* morphs, such that MR expression in the brains of nesting males is three times higher than in sneaker and satellite males (Nugent et al., 2016). In addition, nesting males and satellites have significantly lower levels of circulating CORT than sneakers, and these differences were correlated with differences in neural MR expression, supporting the well-established negative feedback relationship between CORT and its receptors in the brain (Nugent et al., 2016). We also found that MR activity was negatively associated with territorial aggression (Nugent et al., 2016), similar to the situation in plainfin midshipman fish (Arterbery et al., 2010). Further, MR expression in the dorsolateral telencephalon (area Dl), the putative teleost homologue of the mammalian HIP (O’Connell & Hofmann, 2011b), was significantly correlated with courtship behavior in both satellite and nesting males, suggesting an important role for MR signaling in social and sexual behaviors in *S. ocellatus* (Nugent et al., 2016).

In the present study, we aimed to better understand the role of MR signaling in *S. ocellatus* and, more generally, the role of glucocorticoid and AVP signaling in neurogenesis and synaptogenesis in regulating social behavior in the wild. To this end, we designed and intracerebroventricularly (ICV) delivered antisense morpholino nucleotides to knock down MR expression in the brains of *S. ocellatus* nesting males in their natural habitat, followed by quantitative observations of social and reproductive displays. Next, we quantified the expression of genes involved in HPI signaling (GR1, MR, CRF) as well as the immediate-early gene c-Fos, a marker of neural activation in three key nodes of the SDMN associated with stress responsivity and social behavior: the POA, area Dl (the putative teleost homolog of the mammalian HIP), and area Dm (the putative homolog of the mammalian blAMY). To assess the impact of CORT signaling through brain MR receptors on neuroplasticity, we also measured in these brain regions several markers of neuroplasticity (DXC, neurabin, BDNF). In the POA, we also measured the expression of AVP, which regulates HPA/I axis function (Aguilera & Rabadan-Diehl, 2000) and reproductive behavior in *S. ocellatus* (Stiver et al., 2019). Based on our previous research (Nugent et al., 2016) and the broader literature discussed above, we hypothesized that a MR knockdown will increase aggressive and decrease courtship behavior, and therefore reduce successful spawning events. We further hypothesized that these effects would be accompanied by compensatory changes in circulating hormone levels as well as the expression of CRF, AVP, GR1, and MR in the three brain regions under study. Finally, we hypothesized that c-Fos, DXC, neurabin, and BDNF would show increased expression in response to a suppression of MR signaling. This innovative approach allowed us to directly examine the consequences of changes in the HPA/I axis in the wild and identify potential targets of selection among key candidate pathways underlying social behavior and plasticity.

## MATERIALS AND METHODS

### Animals and Injections

All behavioral observations and sample collections were obtained during *S. ocellatus*’s breeding season between mid-May to mid-June 2014 at the University of Liege Marine Laboratory (La Station de Recherches Sous-Marine et Oceanograhic, STARESO), near Calvi, Corsica, France. Reproductively active nests with a territorial nesting male spawning with at least two females, and with a satellite and at least two sneakers and present, were chosen for this experiment. Initial nesting male capture and injections were performed between 10am and 12:30 pm. Nesting males and activity at the focal nest (n=28) were filmed for 10 minutes (see details below). After filming, the nesting male was immediately captured with hand nets, and transported to the lab. Nests were covered with nets during injections to prevent nest takeover by competing males and/or egg predation; nets remained in place until the nesting males were released back to the nest (see below). Within 10 minutes of capture, fish were anesthetized with MS-222 (100 mg/L in seawater for 30 seconds) and placed in a custom immobilization apparatus, wherein fish were submerged in fresh seawater with their heads above water.

Intracerebroventricular (ICV) injections were made into the third ventricle using the third orbital bone as a rostral-caudal landmark and the midline to determine medial-lateral coordinates. The skull was pierced with a 25G needle to a depth of 3.5mm from the top of the head. A Hamilton Neuros Syringe (#7001) with depth control sleeve set to 3.5mm was guided into the third ventricle and 1ul of 400nmol/µl anti-MR or mismatch morpholino (see details below) was injected over 30 seconds. Volume and dose were determined based on previous work in the zebrafish brain (Kizil et al., 2013) and in the neonatal rodent brain (Nugent et al., 2012). Note that in the teleost brain morpholino-mediated knockdown of gene expression is expected to occur within about twelve hours after injection (Kizil & Brand, 2011). Initial sample sizes were n=15 each for treatment and control. The syringe was left in place for an additional 15 seconds post-injection, slowly removed, and a small amount of medical-grade adhesive (VetBond) covered the injection site to prevent leakage. Fish regained their normal equilibrium and locomotor activity after anesthesia within 5 minutes and were delivered back to their nests approximately 25 minutes after initial capture. As one male from each condition failed to return to their nest post-treatment, the study continued from this point with n=14 per treatment. The following afternoon, another 10-minute video of male activity at the nest was filmed. Two days post-injection, a third 10-minute video was taken immediately prior to recapturing injected nesting males with hand nets and collecting their nests to determine emergent larva number and paternity. Fish were euthanized using a lethal dose of MS-222 (2g/L in seawater), blood was collected immediately, followed by rapid decapitation and dissection of brain and gonads. All biological samples were harvested within 20 minutes of final capture. Brains and gonads were stored in RNAlater (Ambion) for later processing (see below). Of the 28 animals who successfully returned to their nest post-treatment, eight were excluded from further analysis for either abandoning their nest post-return (n=3 each, treatment and control) or losing their nests to takeover by another male (n=1 each, treatment and control) prior to final capture. Pre- and post-injection videos were subsequently scored, and the resulting data used in the behavioral analyses described below. The time it took to capture each animal initially and for euthanasia (catch latency) was recorded. All researchers involved in filming and catching the animals, and in scoring the videos, were blind to treatment.

Injection accuracy was determined in a separate cohort of 6 nesting males, which received methylene blue injections into the third ventricle instead of morpholinos. These animals were euthanized approximately 1-minute post injection, and injection accuracy was accessed based on dye perfusion throughout the brain. Four of the six animals had accurate perfusion based on dye injections. The Yale University Institutional Animal Care and Use Committee approved all procedures involving animals (Yale IACUC Protocol number 2014-10908).

### Morpholino Design

Morpholino oligomers provide a potent tool to knock down the expression of candidate genes of interest (Summerton, 2017) in a manner that is much more specific than pharmacological manipulations with receptor antagonists, which often cross-react. Morpholinos comprise oligonucleotides (∼25 base) that are complementary to specific sequences of mRNA of the target gene. Vivo-Morpholinos, which are modified with covalently attached chemical groups to facilitate entry into cells, were custom designed by GeneTools, LLC using proprietary software against a region flanking the start codon of the *S. ocellatus nr3c2* gene (obtained from the *S. ocellatus* transcriptome, Nugent et al., 2016) to block target translation (5’ TTTGGTATCTCTTGGTCTCCATTGC 3’). A 5-base mismatch was used as a control morpholino (5’ TTTaGTATaTCTTaGTCTaCATTaC3’). Blasting this mismatch sequence against the *S. ocellatus* transcriptome returned no results.

### Quantitative real-time PCR

Brains were embedded in TissueTek O.C.T. compound (Sakura Finetek) and sliced on a cryostat in the coronal plane at 300 μm. A 300 μm diameter sample corer tool (Fine Science Tools, Foster City, CA, USA) was used to micro-dissect the three candidate brain regions (POA, area Dl, and area Dm) from the appropriate brain sections, anatomically defined following Maruska et al. (2013). Two microdissected punches (left and right hemisphere) were taken from a single brain slice and stored in DNA/RNA Shield (Zymo Research, Irvine, CA, USA) at −80∘C until further processing. Note that microdissections could not be completed successfully for two samples (n=1 each, treatment and control), which therefore were not included in the analysis below.

We assayed the expression of 7 genes (GR1, MR, neurabin, BDNF, CRF, DXC, c-Fos) in all three brain regions (POA, area Dm, area Dl). AVP was assayed only in the POA, as it is not expressed in areas Dm and Dl (Butler & Maruska, 2020; but see Santiago Rodriguez et al., 2016). RNA was extracted from tissue punches (300µm thick, 300µm diameter) follwing a Proteinase K digestion (at 55°C for 2 hours) using a Quick-RNA MicroPrep kit (Zymo Research) according to the manufacturer’s instructions. cDNA was synthesized using the GoScript Reverse Transcription system (Promega) using random primers and oligo(dT). Primers for region-specific gene expression analysis were designed against sequences in the *S. ocellatus* transcriptome as previously described (Nugent et al., 2016) and are shown in Table 1. Transcript expression was quantified in triplicate for each gene on a ViiA7 Real-time PCR System (Life Technologies) using GoTaq qPCR Master Mix (Promega). Standard curves for each gene were generated from serial dilutions of purified PCR products for each gene. Following the cycling protocol, continuous fluorescence was measured to generate a melting curve from 60°C to 95°C. ViiA7 software automatically generate baseline and threshold values for each gene, and the threshold cycle (Ct) values for each sample were used to determine cDNA quantity. Primer amplification efficiences and relative expresion levels were determined using MCMC.qpcr Bayesian analysis pacakge in R (Matz, Wright & Scott, 2013), with guanosine triphosphate binding protein (GTPbp) and elongation factor 1 (EF1) as control genes. Normalized relative expression values were analyzed by one-way ANOVA across morphs.

**Table 1:**
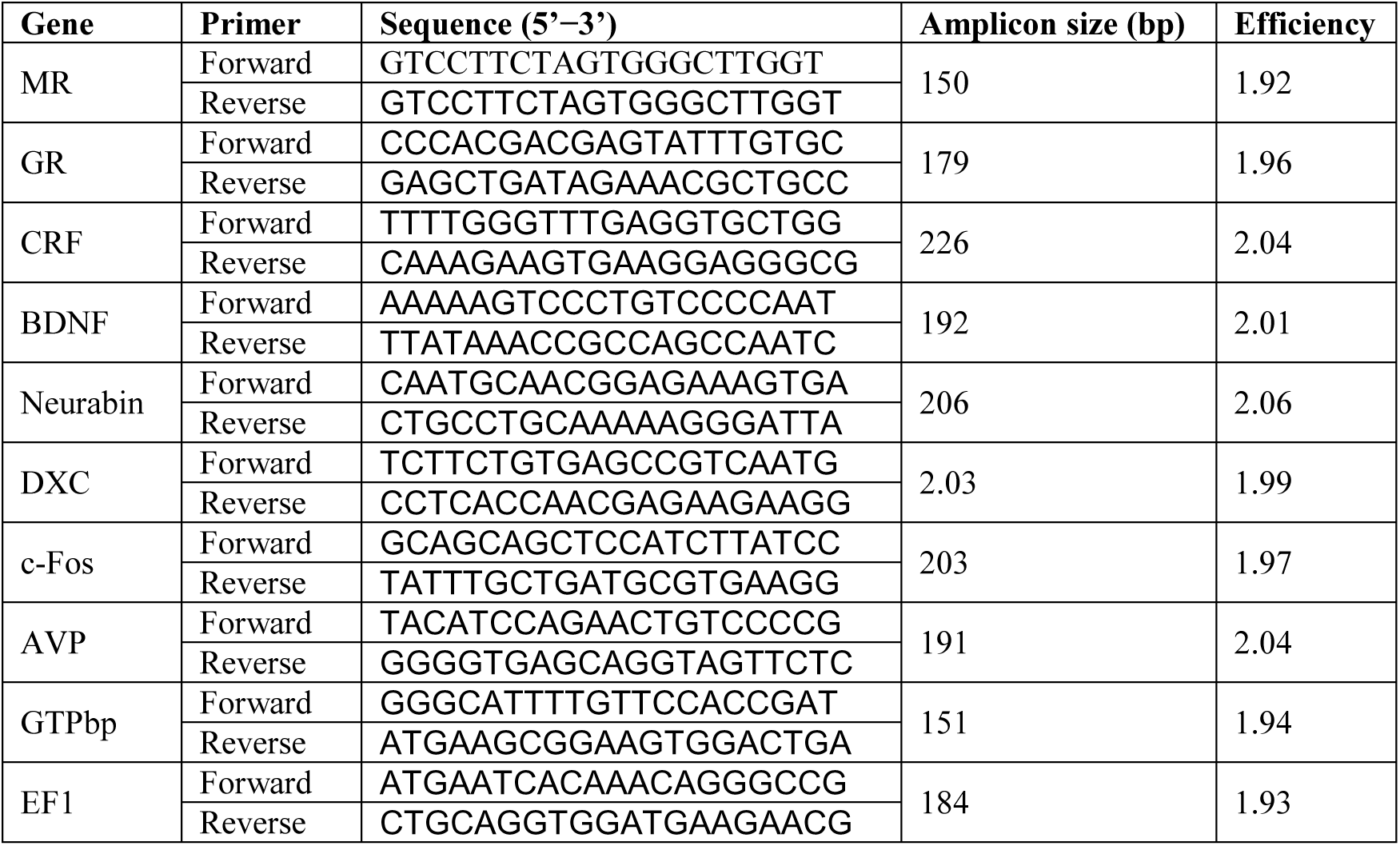
Primer sequences used for quantitative real-time PCR in region-specific analyses

### Hormone Assays

Commercially available enzyme immunoassay (EIA) kits were used to quantify the levels of free plasma CORT (Enzo Life Sciences) and the major teleost androgen, 11-ketotestosterone (11-KT), (Cayman Chemical) as per the manufacturer’s instructions and as previously described (Kidd, Kidd, Hofmann, 2010). Plasma was diluted 1:30 for 11-KT measurement and 1:50 for CORT quantification.

### Behavioral Observation and Analysis

Behavior data was scored from pre-treatment and post-treatment day 2 videos for 10 anti-MR males and 8 mismatch-morpholino males (video file failure prevented behavioral analysis of two mismatch-morpholino males). Nests were filmed for 10 minutes prior to nesting male removal and injection to provide a baseline for comparison. On the day of treatment, nests were checked in the afternoon to assess whether the nesting male had returned to and retained his nest. We then filmed nests for 10 minutes two times after injection: at 24 hours (day 2, post-treatment observation 1) and 48 hours (day 3, post-treatment observation 2) after the initial pre-treatment observation and capture. We later scored behavior on the pre-observation day and day 3 (day 2 post-injection) videos. Day 2 videos were initially taken as insurance of behavioral data due to concern over the potential for nesting male replacement; however, as the majority of males remained in their position until day 3, that second post-treatment observation was selected for scoring and analysis to ensure influence of the injection on behavior. Researchers filming (SMR, SHA and HKK) and scoring (KAS) the behaviors were blind to treatment group. The number of sneakers present at the nest was recorded every minute, and nesting male and satellite proximity to nest was recorded every 30 seconds (coded as “in”: within 10 cm of the nest or closer, “near”: between 10 and 50 cm from the nest, and “far”: further than 50 cm from the nest). It was also noted if the satellite was absent entirely from the nest, as happened with some of the post-injection observations. The total number of females that visited and mated at the nest were recorded, as well as the number of mating bouts events per female. Counts of specific reproductive (courtship; mating bouts), parental (fanning; nest maintenance), and aggressive behaviors (aggression to other nesting males; aggression to subordinate males; aggression to females) performed by the nesting male were recorded. Instances of satellite aggression towards sneakers, and satellite and sneaker submission to the nesting male were documented. Sneak spawns by satellites and sneakers were recorded, as well as the average number of sneakers at nest and whether sneakers and satellites sneak-spawned when the nesting male mated with a female (see Stiver and Alonzo 2013 for more information on behaviors and behavioral scoring procedures).

### Larval Counts and Paternity

Following the final capture of the nesting male, nests (which are made of algae) were collected and secured in individual plankton nets (Fieldmaster 153 μm, 8-inch-diameter Student Plankton Nets). These nets were closed with a cable tie and secured underwater at approximately 3 meters depth using rope and carabiners. Corks were attached to the end of the plankton nets with the sample bottles, such that the sample bottles would float above the main body of the plankton nets. This allowed the larvae (from the nest’s algae secured in the plankton nets) to be collected in the bottle upon hatching, as larval fish of this species swim up after hatching at night (Lejeune 1985). These larval sample bottles were removed and replaced daily. Daily samples were filtered through a Whatman #1 paper filter to collect the larvae into a condensed sample, preserved in 80% ethanol in a 15ml Falcon tube. These samples were later counted individually under a dissecting microscope using a Ward zooplankton counting wheel to quantify the total number of larvae hatching from the nest each day.

### Paternity assignment

We conducted paternity analyses to assign parentage, focusing on collection days 3 and 4, to assure assessment of eggs fertilized after the morpholino injection. For one nest in each condition, no larvae emerged on day 4, and so only day 3 was used. Samples were sent to the University of Arizona Genetics Core (Tucson, AZ, USA) for genotyping. For a detailed description of full DNA extraction, amplification, and scoring methodology see Milazzo et al (2016). Briefly, DNA was extracted using magnetic bead-mediated robotic extraction (Verde Labs Genomic DNA Extraction Chemistry on a Biosprint96 Extraction Robot). All samples were amplified using six microsatellite loci developed for *S. ocellatus* (Soc1017, Soc1063, Soc1109, Soc1198, Soc3121, Soc3200), with modified primer lengths that allowed all six loci to be used in combination in a single PCR in a DNA Engine Tetrad 2 thermal cycler from BioRad set at the following parameters: 94°C (120s); 15 cycles of 94°C (30s), 60–54°C (30s, 60°C on first cycle, decreasing by 0.5°C for each subsequent cycle), 72°C (90s); 23 cycles of 94°C (30s), 54°C (30s), 72°C (90s); 72°C (10 min). Fragment analysis, and visualization and scoring were conducted using an Applied Biosystem 3730 DNA Analyzer, and the standard protocol for GENEMARKER software from Softgenetics. Peaks were visually evaluated and scored by two observers who were blind to sample identity.

We genotyped 25-30 larvae per collection day or the entire sample if the total was less than 25. The average number of larvae genotyped per nest per day was 24. In total, 16 nesting males and 725 larvae from 16 nests (8 per condition) were genotyped; the sample failure rate for larvae was 5%, resulting in 691 larvae that were successfully genotyped. Larvae were included in the analysis only if they could be compared with the putative father at three or more loci (of 691 larvae that were successfully genotyped, five could not be compared to the putative father). We assigned paternity using strict exclusion (see Alonzo & Heckman 2010 for a comparison of different parentage assignment methods using these loci in *S. ocellatus*). To maintain a conservative estimate of nesting male paternity, eggs that had at least one mismatch to the putative father were left unassigned.

### Statistical Analyses

All statistical analyses were carried out in R (version 3.6.1). As described above, we excluded eight samples (nesting males), because they either abandoned or failed to return to their nests (see above), and an additional two samples, because the brain microdissections could not be completed successfully. For all remaining samples (n=18) any missing data (6.57% of all cells) was interpolated by using the mean of the respective measure. This resulted in a dataset with n = 9 MR knockdown and n=9 control animals. The complete dataset is provided in Supplementary Table 1. We initially conducted Student’s t-tests for all individual measures to compare treatment and control groups. Because of the multi-variate nature of the dataset, we conducted Principal Components Analyses (PCAs). We first examined the organism-level data – comprising all available measures of behavior, physiology, and fitness proxies for the differences between post- and pre-treatment (Post – Pre) as well as the post-treatment data only. Next, we conducted PCA on the complete gene expression dataset, comprising all three brain regions examined, followed by a PCA for each brain region separately.

To examine the covariance structure between organism-level measures with gene expression data, we performed a hierarchical clustering analysis of the correlation matrices constructed from organism-level data and the principal component scores obtained from the gene expression analyses, separately for the Post – Pre difference data and the post-treatment only data. Specifically, the Pearson correlation coefficient was calculated for each cell in the matrix (i.e., every measure was correlated against every other measure for all samples). We then performed a hierarchical clustering analysis of the resulting correlation matrix, using average linkage as agglomeration method and correlation as distance metric. Using the R package *pvclust* we estimated the robustness of any resulting clusters by multiscale bootstrap resampling. Clusters for which p < 0.05 are indicated by rectangles (Suzuki and Shimodaira, 2006).

## RESULTS

### MR knockdown does not alter behavior, physiology, or fitness proxies

To test the hypothesis that a knockdown of MR increases aggressive and decreases courtship behavior, and subsequently reduces successful spawning events, we first examined whether the morpholino-mediated MR knockdown in the brain affected any of the behavioral or physiological measures we recorded. There were no significant differences between treatments in changes from baseline (**Post – Pre**) in most of the behavior variables we measured (results not shown). However, it took significantly longer to re-catch MR antisense-treated nesting males following the final behavioral observation compared to mismatch controls (Student’s t-test: t_1,16_ = 2.979, p (uncorrected) = 0.01), often requiring two to three divers to catch antisense-treated males, whereas mismatch-treated males were all caught by a single diver (all divers were blind to treatment, as noted above). Note, there was no different in catch latency when males were caught for treatment (Student’s t-test: t_1,13_ = -0.370, p (uncorrected) = 0.171). We also hypothesized that MR knockdown will cause compensatory changes in circulating hormone levels, such as a decrease in 11-KT and an increase in CORT. We found no significant differences in plasma 11-KT levels across treatment groups (t_1,16_ = 0.819, p (uncorrected) = 0.43). However, nesting males treated with antisense morpholino had indeed higher levels of free plasma CORT compared to mismatch controls (t_1,16_ = 2.266, p (uncorrected) = 0.05). Importantly, CORT levels did not significantly correlate with the time it took to re-catch males (Pearson correlation: r^2^_16_ = 0.17, p (uncorrected) = 0.09). Finally, we hypothesized that MR knockdown might negatively affect fitness measures of nesting males. However, there were no significant differences in the average number of daily larva emerging from nests of MR antisense and mismatch control males (t_17,99_ = -0.091, p = 0.93). In addition, there were no differences in the proportion of NM-sired larva in nests collected from MR antisense and mismatch control treated males (t_1,14_ = 0.150, p = 0.883).

Because of the many behavioral, physiological, and fitness variables measured in this study, we used principal components analysis (PCA) to reduce the dimensionality of our dataset. When we investigated the changes in organism-level attributes from before to after treatment (**Post – Pre**), neither principal component (PC) 1 (which explained 31.09% of the variance in the data; Student’s t-test: t_1,16_ = 1.4499, p = 0.17) nor PC 2 (12.29%; t_1,16_ = 0.812, p = 0.43) showed any significant clustering by treatment (Supplementary Figure 1). All PC loadings are provided in Supplementary Table 2. Similarly, there was no difference between antisense- and control-treated animals in the **Post-Treatment** dataset, i.e., the organism-level attributes measured after treatment (Figure 1): Neither PC 1 (which explained 38.79% of the variance in the data; t_1,16_=1.217, p=0.24) nor PC 2 (14.17%; t_1,16_=0.302, p=0.77) showed any significant separation by treatment. All PC loadings are provided in Supplementary Table 3. Taken together, while there is considerable variance in our data, we did not find any evidence that a knockdown of MR expression directly affects any of the behavioral or physiological variables or fitness proxies that we measured, with the exception of increased CORT levels in MR knockdown animals.

**Figure 1:**
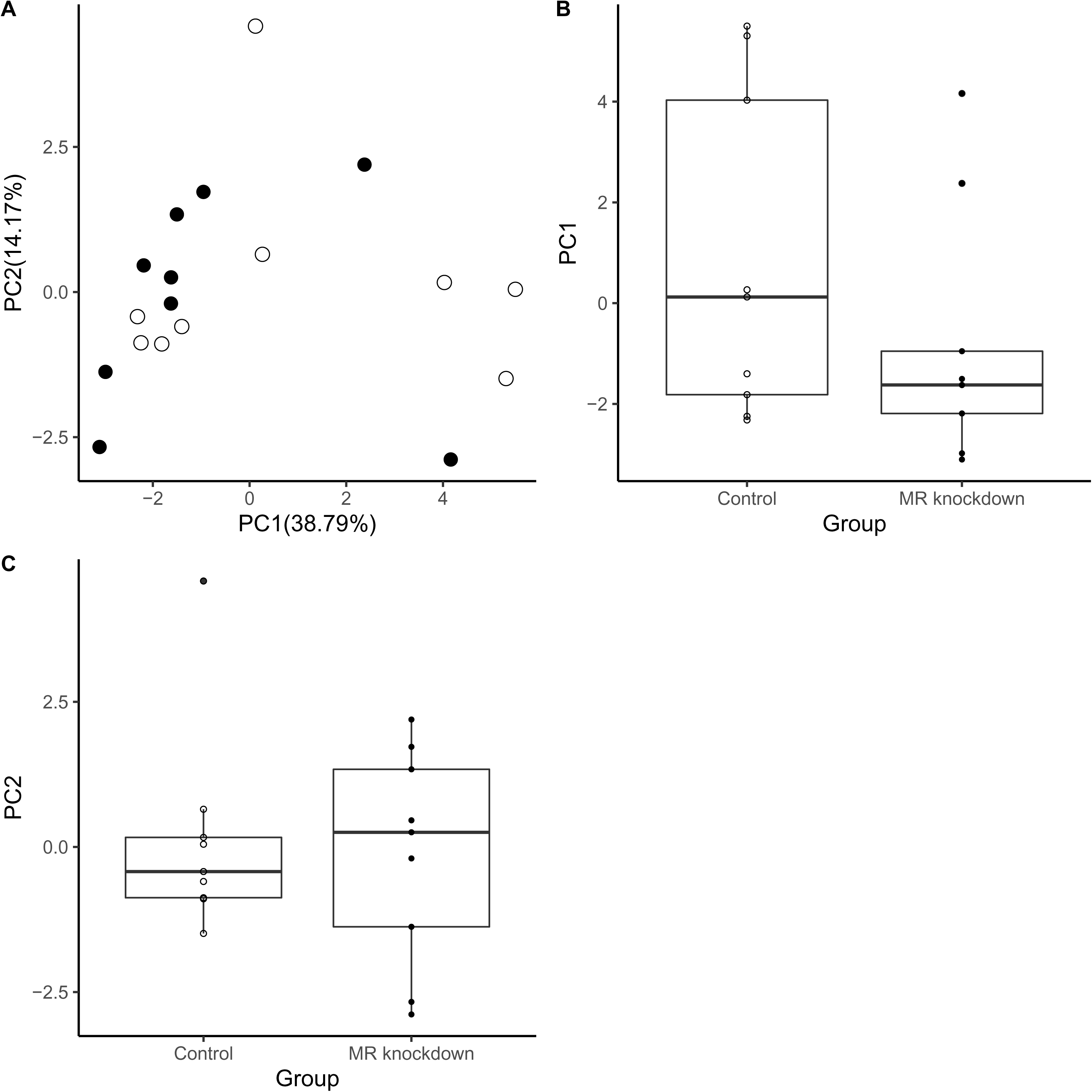
Principal component analysis of all organism-level traits measured at the end of the experiment (**Post Treatment**). **(A)** Scatter plot showing how MR knockdown (filled circles) and control animals (empty circles) are distributed in variance space along PC1 and PC2. **(B)** Box plots indicating that treatment and control did not separate along the axes of PC1 or PC 2.

### MR knockdown affects gene expression in a brain-region-specific manner

Even though the morpholino-mediated knockdown of the brain MR did not appear to have readily observable behavioral or reproductive effects, we did observe significantly increased levels of circulating CORT in MR morpholino-treated males. We had hypothesized that MR knockdown would be accompanied by such compensatory changes, including also concomitant changes in the expression of CRF, AVP, GR1, and MR in area Dl, area Dm, and the POA. We also hypothesized that molecular markers of neural activity, neurogenesis, and synaptogenesis would show characteristic changes in response to a suppression of MR signaling. To test these hypotheses, we first compared the expression levels for each gene in each brain region directly and identified numerous significant differences (results not shown). Even though these results suggested robust effects of MR knockdown on brain gene expression, due to the number of genes assayed, some of these differences were no longer significant once we accounted for multiple hypothesis testing. To examine whether gene expression varied by treatment and/or brain region we therefore performed several different PCAs as a way to reduce the dimensionality of the data.

Because it is well known that gene expression varies considerably across brain regions (Lein et al., 2007), which needs to be taken into account for data analysis, we first examined the expression data for all the genes that were measured in all three brain regions (i.e., GR1, MR, neurabin, BDNF, CRF, DXC, c-Fos). PC1 and PC2 together explained 69.96% of the variance (PC1: 52.06%; PC2: 17.9%; Supplementary Figure 2A) and significantly separated the three brain regions (PC1 ANOVA: F_2,51_ = 367.1; p = 2.2x10^-16^; PC2: F_2,51_ = 67.08; p = 5.3x10^-15^), as expected (Supplementary Figure 2B,C). Tukey *post hoc* tests further demonstrated that gene expression profiles of the POA, area Dl, and area Dm all differed significantly from each other for both PC1 and PC2 (Table 2). All PC loadings are provided in Supplementary Table 4).

**Table 2:** Results of Tukey post hoc tests examining the first two principal components (PC1 and PC2) resulting from the principal components analysis of gene expression variation across the preoptic area (POA), area Dl, and area Dm. (Degrees of freedom = 1,16)

| Comparison | PC1 |  | PC2 |  |
| --- | --- | --- | --- | --- |
|  | test statistic (d.f.) | p-value | test statistic (d.f.) | p-value |
| POA vs. DI | -0.899593 | $4.5 \times 10^{-6}$ | 0.764596 | 0 |
| POA vs. Dm | -4.2500297 | 0 | 2.272345 | $1.01 \times 10^{-3}$ |
| DI vs. Dm | 3.3504366 | 0 | 1.507749 | 0 |

A closer inspection of Supplementary Figure 2A suggested that gene expression patterns not only differ by brain region but also between MR morpholino-treated and control animals in each of the three SDMN nodes. To test this observation we conducted a PCA for each brain region separately. These analyses included expression levels for seven genes (GR, MR, neurabin, BDNF, CRF, DXC, c-Fos) for area Dl and area Dm and all these genes plus AVP for the POA. **POA** gene expression profiles clustered significantly along both PC1 (which explained 44.75% of the variance in the data; Student’s t-test: t_1,16_ = 3.455, p = 3.77x10^-3^), largely driven by BDNF, DXC, and CRF; and PC2 (19.33%; t_1,16_ = -3.159, p = 6.98x10^-3^), driven by GR1, c-Fos, AVP (Figure 2; PC loadings are provided in Supplementary Table 5). **Area Dl** gene expression also clustered significantly by treatment along PC1 (48.7%; t_1,16_ = -5.572, p = 4.28x10^-5^), largely driven by GR1, MR, neurabin, DXC, but not along PC2 (18.03%; t_1,16_ = 0.743, p = 0.47) (Figure 3; PC loadings are provided in Supplementary Table 6). Finally, gene expression profiles in **Area Dm** clustered by treatment only along PC2 (13.15%; t_1,16_ = 3.586, p = 2.58x10^-3^), largely driven by GR1. The variance contained in PC 1 (58.81%) was not explained by treatment (t_1,16_ = 1.302, p = 0.21) (Figure 4; PC loadings are provided in Supplementary Table 7). In sum, although MR morpholino treatment had few significant effects on behavior, physiology, or fitness measures (see above), we found considerable differences in gene expression patterns between MR knockdown and control animals in all three brain regions examined, largely (but not exclusively) driven by genes involved in glucocorticoid signaling.

**Figure 2:**
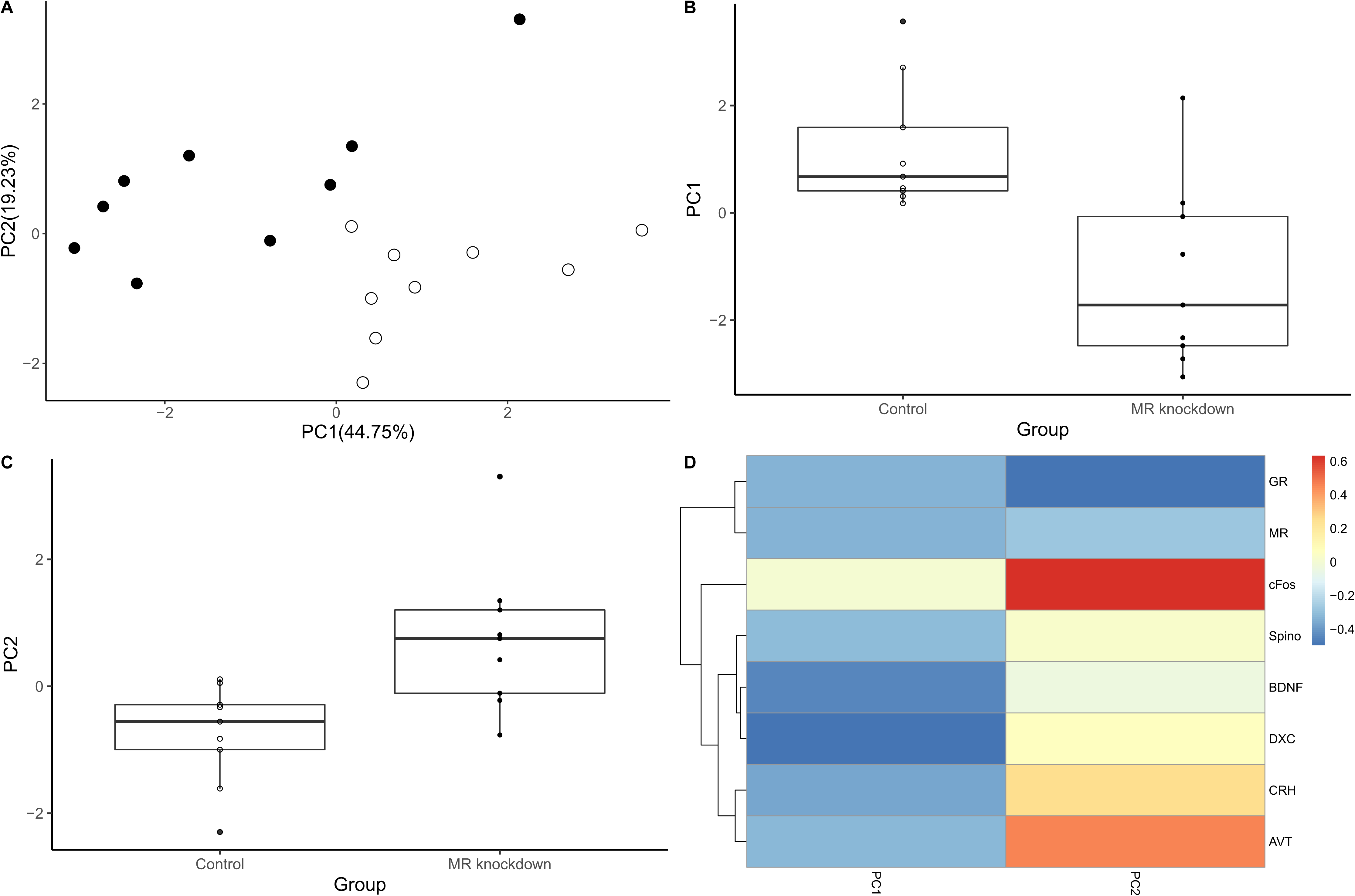
Principal component analysis of the expression data for all the genes assayed in the POA (i.e., GR1, MR, neurabin, BDNF, CRF, DXC, c-Fos, AVP). **(A)** Scatter plot showing how POA gene expression profiles of MR knockdown (filed circles) and control animals (empty circles) are distributed in variance space along PC1 and PC2. Box plots indicating that POA gene expression differed significantly between treatment groups for both **(B)** PC1 and **(C)** PC 2. **(D)** Clustered heatmap that shows how the various factors (gene expression levels) load on each PC. See text for an explanation of gene symbols.

**Figure 3:**
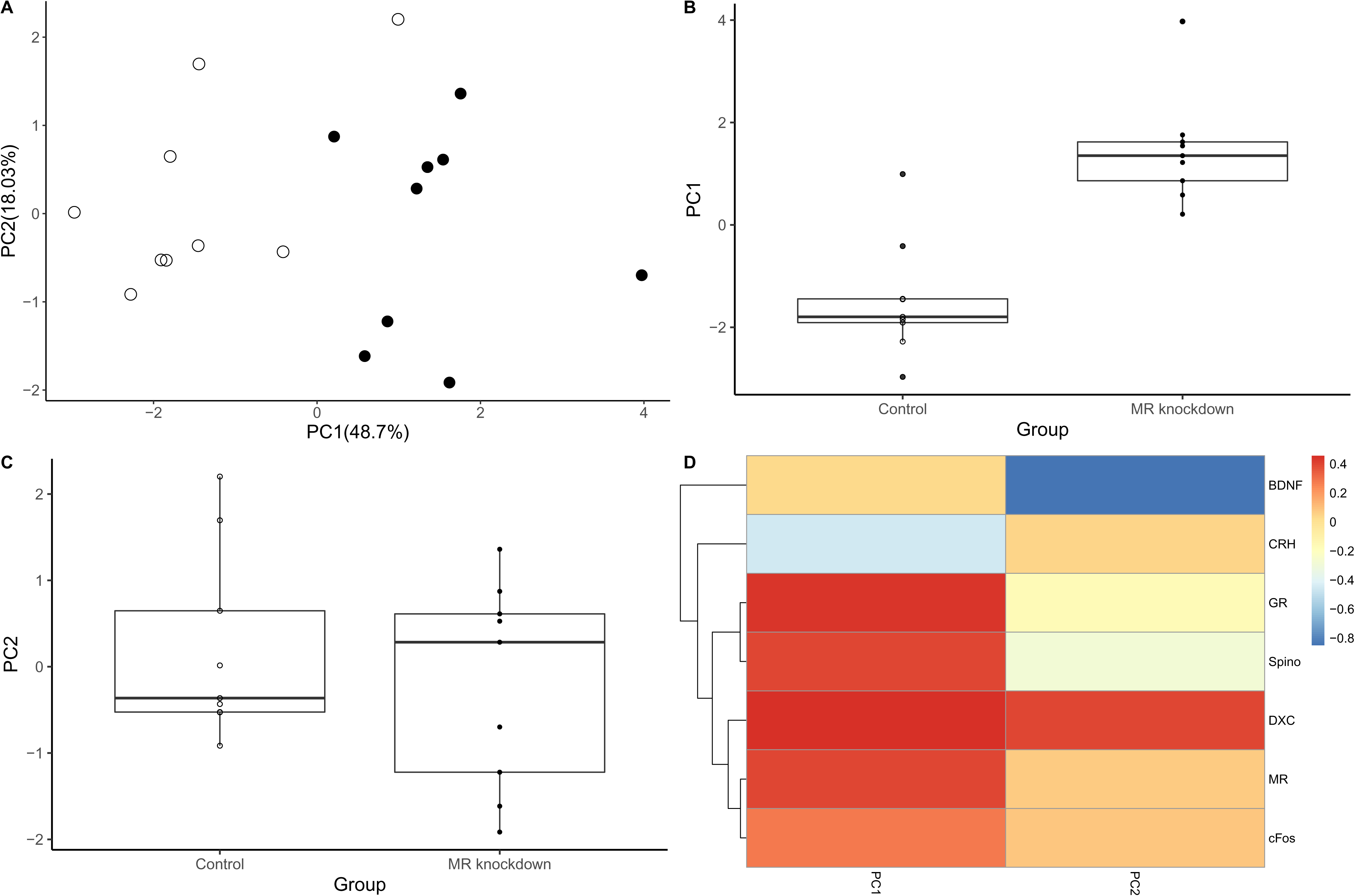
Principal component analysis of the expression data for all the genes assayed in area Dl, the putative teleost homolog of the mammalian hippocampus (i.e., GR1, MR, neurabin, BDNF, CRF, DXC, c-Fos). **(A)** Scatter plot showing how area Dl gene expression profiles of MR knockdown (filled circles) and control animals (empty circles) are distributed in variance space along PC1 and PC2. Box plots indicating that area Dl gene expression **(B)** differed significantly between treatment groups only for PC1 but not **(C)** PC2. **(C)** Clustered heatmap that shows how the various factors (gene expression levels) load on each PC. See text for an explanation of gene symbols.

**Figure 4:**
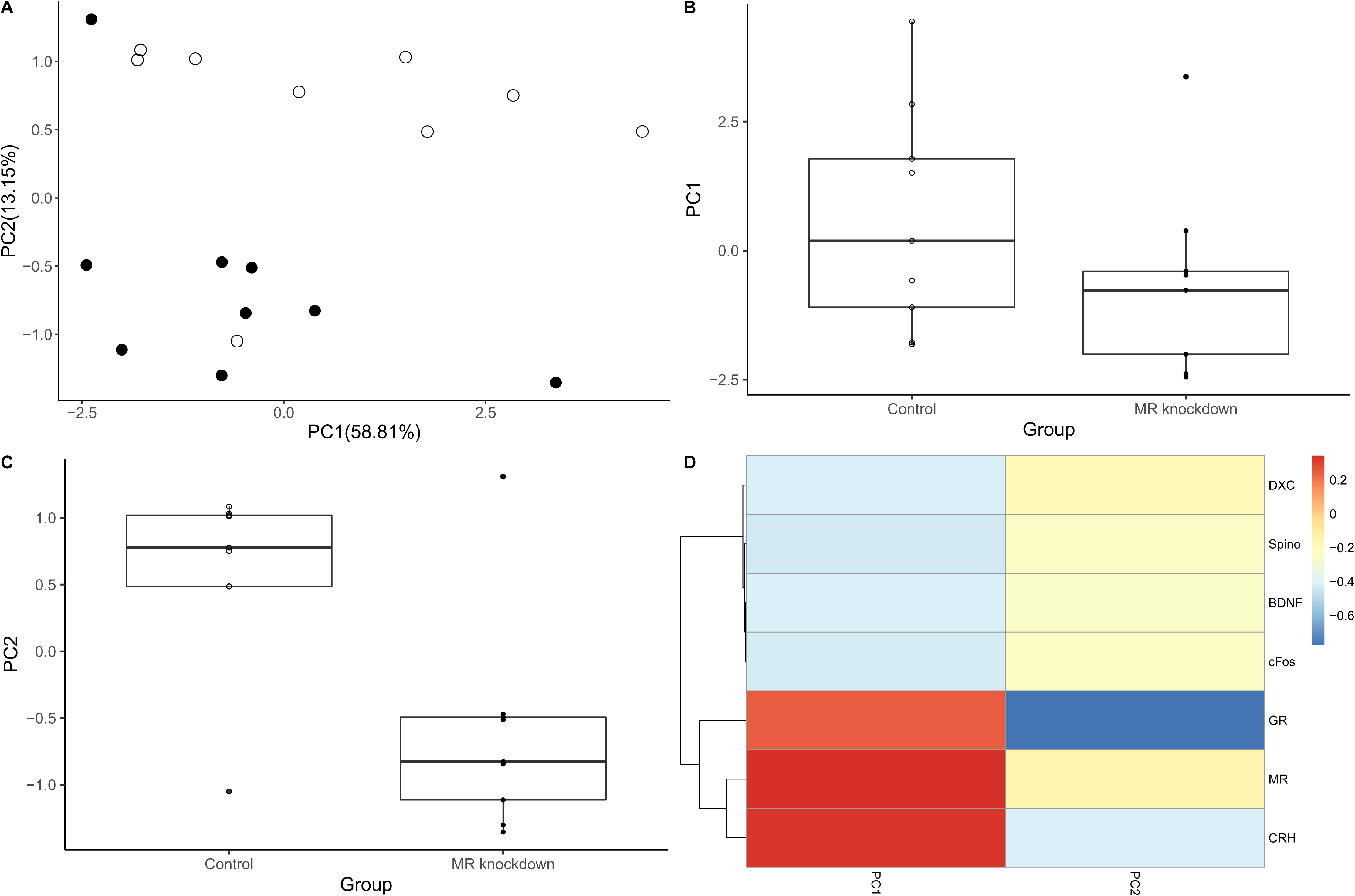
Principal component analysis of the expression data for all the genes assayed in area Dm, the putative teleost homolog of the mammalian basolateral amygdala (i.e., GR1, MR, neurabin, BDNF, CRF, DXC, c-Fos). **(A)** Scatter plot showing how area Dm gene expression profiles of MR knockdown (filled circles) and control animals (empty circles) are distributed in variance space along PC1 and PC2. Box plots indicating that area Dm gene expression **(B)** did not differ significantly between treatment groups for PC1, but **(C)** did so for PC2. **(C)** Clustered heatmap that shows how the various factors (gene expression levels) load on each PC. See text for an explanation of gene symbols.

### Co-variance analysis reveals how MR knockdown affects phenotypic integration

The considerable variation in neural gene expression induced by MR knockdown makes it possible to uncover how brain gene expression, hormone levels, behavior, and fitness proxies are integrated, and can even suggest functional relationships. Not surprisingly, we found several strong gene-behavior and hormone-behavior relationships for the **Post – Pre**-treatment data (Supplementary Table 8) and the **Post-Treatment** data (Supplementary Table 9). Many of these correlations are suggestive of a robust integration of brain region-specific gene expression profiles with behavior, physiology, and fitness proxies of nesting males, though only a subset would remain significant after controlling for multiple hypothesis testing. We instead examined these dataset by creating co-variance matrices for all organism-level measures, as well as the first two PCs of the brain region-specific gene expression PCAs (which together explained between 64% and 72% of the total variance, depending on the brain region). We then identified significant (i.e., p ≤ 0.05) clusters by bootstrapping analysis. We first did this for the treatment minus baseline (**Post – Pre**) differences, which showed little robust clustering of organism-level attributes with brain gene expression, as expected (Supplementary Figure 3). The interesting exception was a robust association between Aggression towards Nesting Males and Sneak Attempts and PC2 of the POA gene expression analysis (which was most strongly driven by the expression of GR1, c-Fos, AVP, see above).

Because brain gene expression profiles are more likely to be associated with the behavioral and physiological measures obtained immediately before euthanasia, we repeated this analysis for **Post-Treatment** organism-level data in conjunction with the first two PCs of the gene expression PCAs presented above. Bootstrapping analysis identified several significant covariance clusters, where numerous organism-level attributes robustly correlated with each other as well as three clusters where such attributes were significantly associated with brain gene expression (Figure 5). Specifically, circulating CORT levels were significantly associated with PC1 of area Dl (putative HIP homolog; largely driven by GR1, MR, neurabin, DXC) and PC2 of the POA gene expression analyses (largely driven by c-Fos, GR1, AVP). Another cluster comprised two reproductive measures (Gonad mass and the proportion of offspring sired by the Nesting Male) and PC2 of area Dl (putative hippocampus homolog) gene expression analysis (strongly driven by BDNF and DXC). Finally, a small cluster contained Aggression to NMs and PC1 of the Dm expression analysis (which was strongly driven by GR1, MR, and CRF). Importantly, most of these gene expression PCs significantly separated MR knockdown animals from controls (see above). Also of note was the robust correlation between circulating 11-KT levels and parental behavior (Fanning). Taken together, knockdown of brain MR expression uncovered strong phenotypic integration between organism-level attributes and brain gene expression patterns in nesting males of *S. ocellatus*.

**Figure 5:**
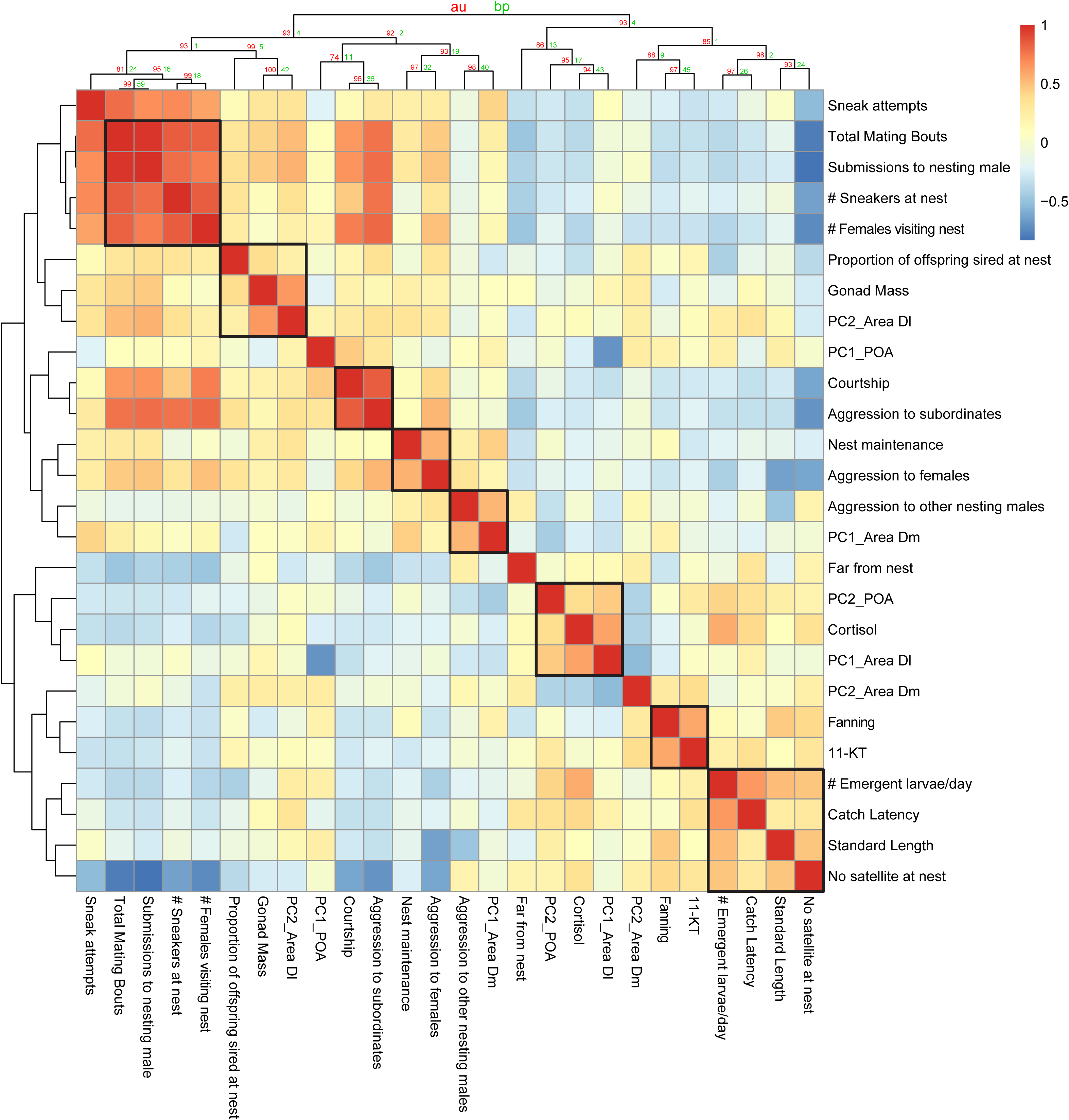
Clustered heatmap of the co-variance matrix for all organism-level attributes measured at the end of the experiment (**Post-Treatment**) along with the first two PCs of each brain region-specific gene expression PCA (shown in Figures 3 to 5). Bootstrapping analysis identified several significant covariance clusters, where organism-level attributes robustly correlate with each other as well as several clusters where such attributes associated with brain gene expression. Significant correlation clusters are indicated by black rectangles.

## DISCUSSION

The study of the neuroendocrine pathways that mediate social behavior in vertebrates offers an opportunity to understand the proximate mechanisms underlying the evolution of complex reproductive behaviors, such as paternal care and alternative reproductive tactics. Such insights are necessary if we want to understand how variation in mechanisms shape behavioral variation, which in turn can affect an animal’s fitness (Hofmann et al., 2014; Rittschof & Robinson, 2014). In the present study, we quantified gene expression patterns in brain regions associated with stress responsivity, social behavior, and cognition to understand the role of MR in these functions in wild, naturally-behaving ocellated wrasse, *S. ocellatus*, nesting males. To our knowledge, our study is the first to use an antisense morpholino to knock down the expression of a candidate gene in a wild vertebrate in its natural environment. Specifically, we suppressed the expression of MR in the brain of nesting males, who engage in courtship and paternal care, and aggressively defend their nest against sneak spawning attempts by satellite and sneaker males. We predicted that MR knockdown would increase aggressive displays by nesting males and, in turn, decrease successful reproduction. We also predicted that MR knockdown would result in compensatory changes in CORT levels and expression of genes involved in glucocorticoid signaling.

### The role of brain MR in regulating behavior, physiology, and fitness

MR knockdown did not affect most of the behavioral or physiological variables we measured nor did it affect either estimate of reproductive success (fitness proxies) we used. However, compared with controls, MR knockdown animals required significantly more time and effort to catch (i.e., Catch Latency), even two days after treatment, and they had significantly higher circulating CORT levels. As there was no significant correlation between the time to catch an animal and circulating levels of CORT, this suggests that CORT levels were not elevated by extended catching times and that instead MR is indeed a CORT receptor in teleosts and, more generally, that MR is important for negative feedback regulation of the HPA/I axis, as is well established in mammals (Joëls et al., 2008). The significant increase in Catch Latency in MR knockdown animals could be due to an enhanced memory of prior catch, or an enhanced fear response associated with a nearby diver. In addition, this shift in behavior could be associated with the MR knockdown males being more risk avoidant, as they had to be willing to leave their nests in order to delay capture.

Assuming MR knockdown did have its intended effects on the brain and CORT physiology (see also next section), we next consider why we did not see more direct effects on behavior and/or fitness proxies. Observing significant effects of pharmacological perturbations in wild animals observed in their natural environment is always complicated by the many biotic and abiotic factors that cause variation in phenotypes of interest that might mask the relatively subtle changes of behavior we predicted. For example, observation times in terms of both duration and time of day are necessarily limited when dealing with an animal that lives underwater. Furthermore, we were not in a position to control the length of the reproductive cycle, nor how long specific males had been in their spawning phase (total cycle length and the length of each portion – building, spawning, and fanning – varies within and among males, Lejeune 1985, Alonzo 2004). It is thus possible that, due to naturally occurring variation among nests, some males were beginning to transition to parental care at the time of injection (a phase where males are quite variable in the time spent engaging in parental care and at the nest; Hellmann et al. in prep). For example, while there was considerable variation among males in how much time they spent away from the nest, this could not be associated with treatment. It is possible that this natural variation amongst nesting males in their reproductive state interfered with our ability to detect statistically significant effects of MR knockdown on male behavior and reproductive success. Further study of the behavioral and fitness effects of MR and associated pathway is certainly warranted in this species and in teleosts in general.

### MR knockdown affects gene expression in a brain-region-specific manner

We found that the considerable variation in organism-level attributes was underpinned by robust brain gene expression differences between MR knockdown and control animals, in support of our original hypothesis. Specifically, our results show that MR signaling underlies the variation in neural activity (c-Fos), glucocorticoid system regulation, neurogenesis (DXC), synaptogenesis (neurabin), neuronal differentiation (BDNF), and learning & memory pathways (BDNF) in a brain region-specific manner. The considerable differences in gene expression patterns between MR knockdown and control animals in all three brain regions examined are largely (but not exclusively) driven by genes involved in glucocorticoid signaling.

In the **POA** most of the genes we examined are upregulated in antisense treated animals, suggesting enhanced cellular activation and plasticity in this region in response to MR reduction. Specifically, expression of the immediate-early gene c-Fos (a marker of neural activity: Morgan & Curran, 1991) is increased in MR knockdown males and loads strongly on PC2 of the POA PCA, along with GR1 and AVP, which are both important in HPA/I axis activation (Aguilera & Rabadan-Diehl, 2000; Joëls et al., 2008; Gesto et al., 2014). In addition, the high CRF expression in MR knockdown males might be driving the increase in circulating CORT we observed. In concert with CRF, preoptic BDNF expression is also increased, possibly as a consequence of increased CORT levels, as in mammals (Smith et al., 1995). BDNF, in turn, has been shown to increase CRF levels in mammals (Toriya et al., 2010), and this might be happening here as well. Finally, we also find increased DXC expression in MR knockdown males (strongly loading on POA PC1), suggesting that MR signaling also regulates neurogenesis in the POA.

Our gene expression data provide additional molecular evidence that **area Dl** (lateral pallium) is indeed the putative (partial) homolog of the mammalian HIP (O’Connell & Hofmann, 2011b). Our finding that the expression of GR1, MR, and CRF load most strongly on PC1 (with increased expression in MR knockdown males) is consistent with the notion that area Dl is a main feedback point for glucocorticoid signaling from the HPA/I axis, a relationship that is well established in the mammalian HIP (Kim et al., 2015). We also find DXC to be regulated inversely to these glucocorticoid signaling genes, possibly indicating reduced neurogenesis in area Dl as a consequence of MR knockdown and a compensatory increase in glucocorticoid signaling. Neurabin expression in area Dl may behave in a similar manner. These findings are consistent with the situation in the mammalian HIP, where both neurogenesis and synaptogenesis (such as formation of dendritic spines) are known to be inhibited by stress axis activation (McEwen, 2017).

The mammalian basolateral (pallial) amygdala has been implicated in fear, memory consolidation, and social recognition (Duvarci & Paré, 2007; Wang et al., 2014). We found that in **area Dm** (medial pallium), the putative teleost homolog of this brain region (O’Connell & Hofmann, 2011b), most of the variation in gene expression patterns did not separate MR knockdown animals from controls. However, PC2, which still explained 13.5% of the total variance, significantly differentiated the two experimental groups, most strongly driven by GR1 expression. We also observed an increase in neuronal activity in this brain region with MR knockdown, as indicated by increased c-Fos expression. Just like the blAMY of mammals, the teleost Dm is involved in fear responses as well as emotional learning (Portavella et al., 2002). Thus, the increased activity in this region could be solely in response to being recaptured or re-observed following initial capture.

### Integration across level of biological organization reveals how MR signaling mediates behavior, physiology, and fitness

An intimate understanding of variation in the physiological and neural mechanisms mediates variation in behavior can help uncover both the evolutionary history of the behavior under study as well as the underlying targets of selection (Phelps et al., 2010; Hofmann et al., 2014). By integrating organism-level and gene expression data within a covariance analysis framework we were able to discover robust relationships between behavioral displays, hormone levels, and brain region-specific molecular pathways that confirm the central role played by MR signaling in the regulation of complex social behavior. Looking at the changes between before and after treatment (**Post – Pre**) we found little robust clustering, which is expected, given that gene expression patterns are unlikely to integrate changes that have occurred over several days but instead be reflective of the behavioral activity before the animals were euthanized. Nevertheless, given the fundamental role of the POA in regulating male aggression across vertebrates (Dominguez & Hull, 2005), it was reassuring to find a strong association between aggressive displays towards other nesting males and PC2 of the POA gene expression PCA, which significantly separated MR knockdown from control animals by variation in the expression of GR1, c-Fos (a neural activity marker), and AVP (which is well known to regulate aggressive behavior in other teleosts, e.g. Greenwood et al., 2008).

When we examined the **Post-Treatment** data, we found clusters of robustly correlated behaviors, which was not surprising. More interestingly, though, we found a strong relationship between CORT levels and area Dl PC1 (most strongly loaded by GR1, MR, neurabin, DXC), which significantly separated MR knockdown animals from controls. This cluster corroborates the results discussed above and suggests differential activation of glucocorticoid signaling and neural plasticity depending on treatment, reminiscent of the classic feedback regulation of hippocampal function by the HPA axis that is well known in mammals, including altered neuroplasticity (McEwen, 2017). Just as exciting, the proportion of offspring sired by nesting males (a proxy of fitness) and their reproductive status (GSI) clustered with PC2 of area Dl (strongly loaded by BDNF), suggesting that hippocampal neuroplasticity mediates evolutionary fitness and thus represents a target of selection. The strong association of aggressive displays toward nesting males and PC1 of area Dm (strongly loaded by GR1, MR, and CRF) provides additional evidence that the putative blAMY homolog of teleosts regulates affective processes via glucocorticoid signaling, similar to the situation found in mammals (Duvarci & Paré, 2007). Finally, the strong association of 11-KT levels with fanning behavior (which directs oxygen-rich water towards the offspring) suggest a role of androgens in regulating paternal behavior, as has been demonstrated in other teleosts (Pradhan et al., 2014; DeAngelis et al., 2018). In sum, our integrative analyses revealed strong phenotypic integration of behavioral and fitness traits with hormone levels and brain region-specific patterns of glucocorticoid signaling and neuroplasticity. Future research should examine to which extent these neuromolecular mechanisms serve as targets of selection.

## CONCLUSIONS

Neurobiologists and evolutionary biologists increasingly focus on the neuroendocrine pathways that underlie complex behavioral phenotypes, including courtship, nest defense, and parental care, because they provide an intuitive connection between proximate mechanisms of behavior and their evolutionary consequences, and as such are key to explaining the diverse reproductive behaviors of animals (Hofmann et al., 2014; Rittschof & Robinson, 2014). However, to date most studies have, by necessity, focused on model species or captive populations, which can paint an incomplete or skewed picture of the true contribution of a given mechanism to the evolution of complex behaviors in nature. It is essential that we also study patterns of behavior and fitness under natural conditions in the wild, if we wish to move beyond the “phenotypic gambit” (Grafen, 1984) and expand our understanding of the evolutionary forces that have shaped these behaviors and the mechanisms underlying them. In the present study, we experimentally manipulated a central signaling pathway in a wild population. We measured a suite of physiological and behavioral responses, and quantified a robust proxy of fitness (number of larvae hatched), thereby addressing the relative contributions of the MR signal pathway to reproductive behavior and fitness of nesting males. While this necessitated the introduction of uncontrolled variables into our study design (such as the timing of the nest cycle), we were able to show that the complex mating and paternal care behavior in our focal species is associated with the MR pathway, consistent with our knowledge of caring behavior in other vertebrates (Krause et al., 2015). In the future, measuring gene expression profiles (transcriptomes) could provide a way of measuring phenotypic metrics in natural settings that are less susceptible to environmental and measurement noise. This novel field of quantifying behavioral ecological transcriptomics in wild population will undoubtedly allow for new discoveries of the role of these conserved neuroendocrine pathways in the evolution of diverse social and reproductive behaviors.

## Supporting information

Supplemental Tables - combined

## ACKNOWLEDGEMENTS

We thank Natascia Tamburello for assistance in the field as well as Dr. Pierre Lejeune, Alexandre Volpon and the entire staff at the Station de Recherches Sous-Marines et Oceanographique (STARESO). We thank Emma Wilson for counting the larval samples and preparing them for parentage analysis and the Arizona Research Laboratory at the University of Arizona Genetics Core for assistance with microsatellite analyses, and Raymond Mungo for consulting on the statistical analyses. We also thank Emily Lessig and Tessa Solomon-Lane for comments on earlier drafts of the manuscript and members of the Hofmann laboratory for discussion. This research and Suzanne Alonzo were supported by the National Science Foundation (under Grant Numbers IOS-0950472 and IOS-1655297) and Yale University. Holly Kindsvater was supported by a National Science Foundation fellowship (DBI 13-05929). Bridget Nugent was supported by a Gaylord Donnelly Environmental Postdoctoral Fellowship from the Institute for Biospheric Studies at Yale University. Kelly Stiver was supported by SCSU and CSU-AAUP Faculty Development Grants. Hans Hofmann was supported by the National Science Foundation (under Grant Numbers IOS-1354942 and IOS-1501704).

## Supplementary Figures

**Supplementary Figure 1:** Principal component analysis of changes in all organism-level attributes between before to after treatment (**Post – Pre**). **(A)** Scatter plot showing how MR knockdown (filled circles) and control animals (empty circles) are distributed in variance space along PC1 and PC2. Box plots indicating that treatment and control did not separate along either **(B)** PC1 or **(C)** PC 2.

**Supplementary Figure 2:** Principal component analysis of the expression data for all the genes (i.e., GR1, MR, neurabin, BDNF, CRF, DXC, c-Fos) assayed in all three brain regions. **(A)** Scatter plot showing how gene expression profiles for the three brain regions (squares: POA; circles: area Dl; triangles: area Dm) are distributed in principal components space along PC1 and PC2. MR knockdown and control animals are indicated as filled and empty circles, respectively. Box plots indicating that the three brain regions significantly differ along the axes of **(B)** PC1 and **(C)** PC 2, as determined by ANOVA. Letters indicate significant differences between brain regions according to Tukey *post hoc* tests.

**Supplementary Figure 3:** Clustered heatmap of the co-variance matrix of the changes in all organism-level attributes between before to after treatment (**Post – Pre**) as well as with the first two PCs of each brain region-specific gene expression PCA (shown in Figures 3 to 5). Bootstrapping analysis identified few significant covariance clusters (indicated by black rectangles).

