## Supplementary figures and images for "Morpholino-mediated knockdown of the brain mineralocorticoid receptor affects glucocorticoid signaling and neuroplasticity in wild ocellated wrasse (*Symphodus ocellatus*)"

### Supplemental Figure 1

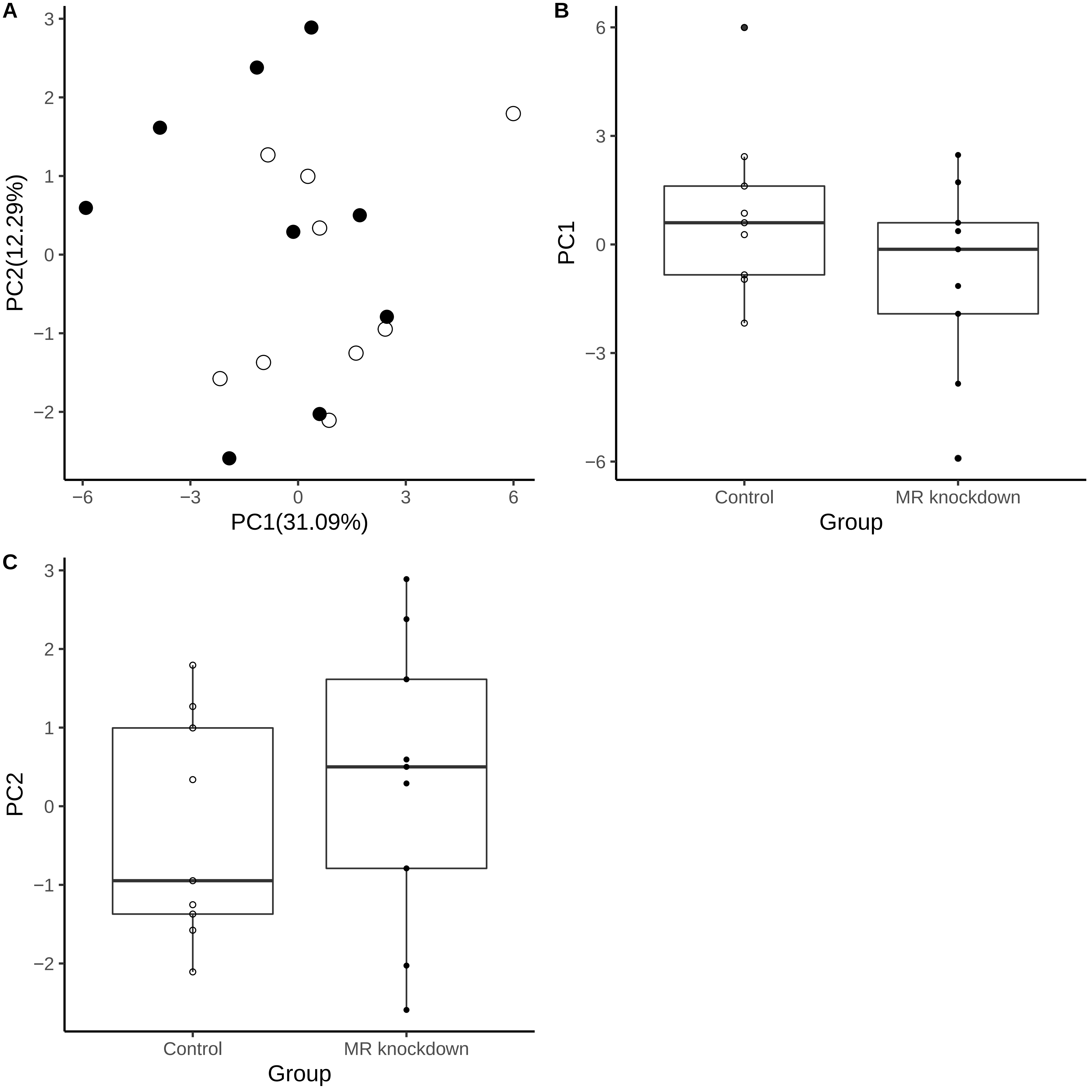

### Supplemental Figure 2

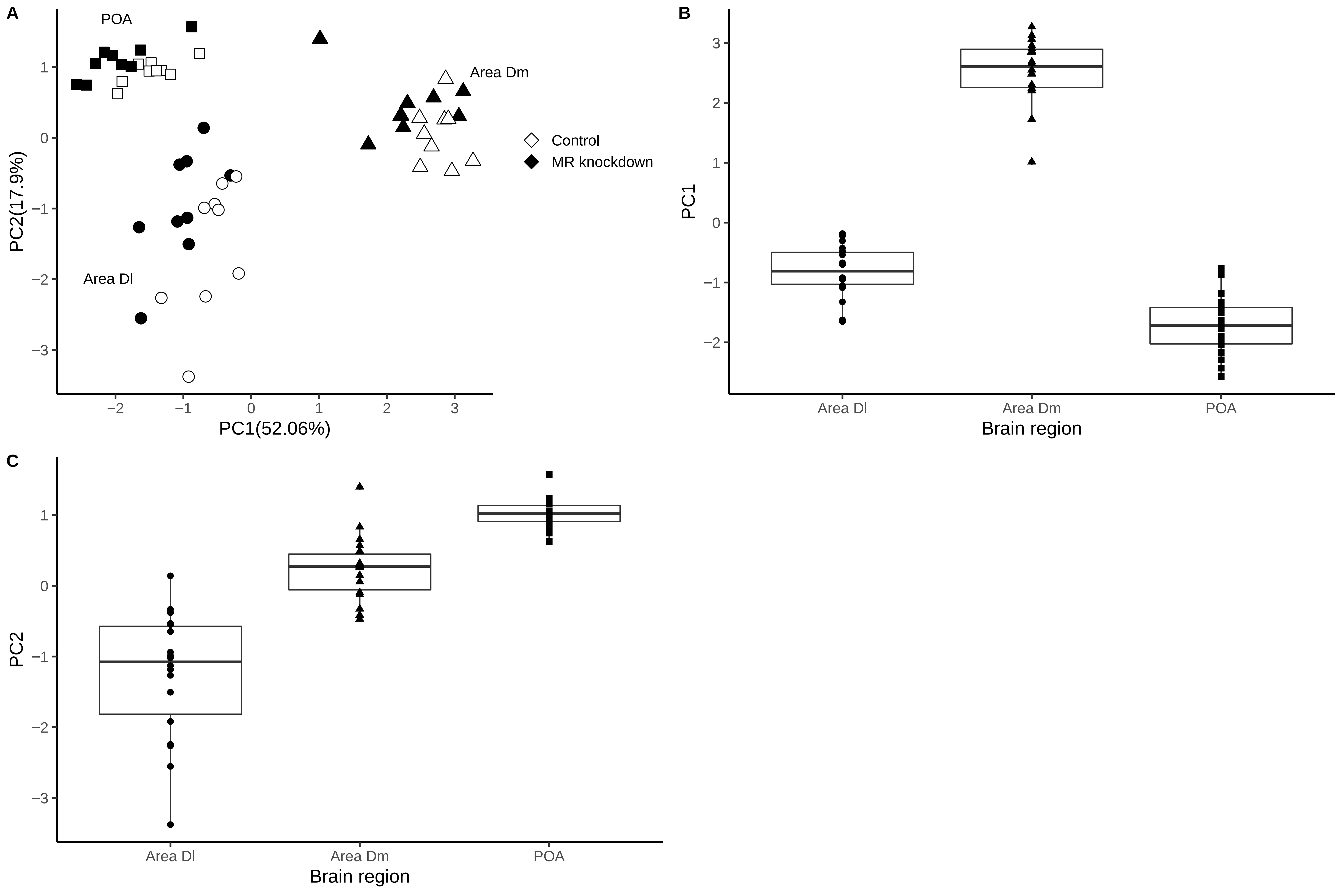

### Supplemental Figure 3

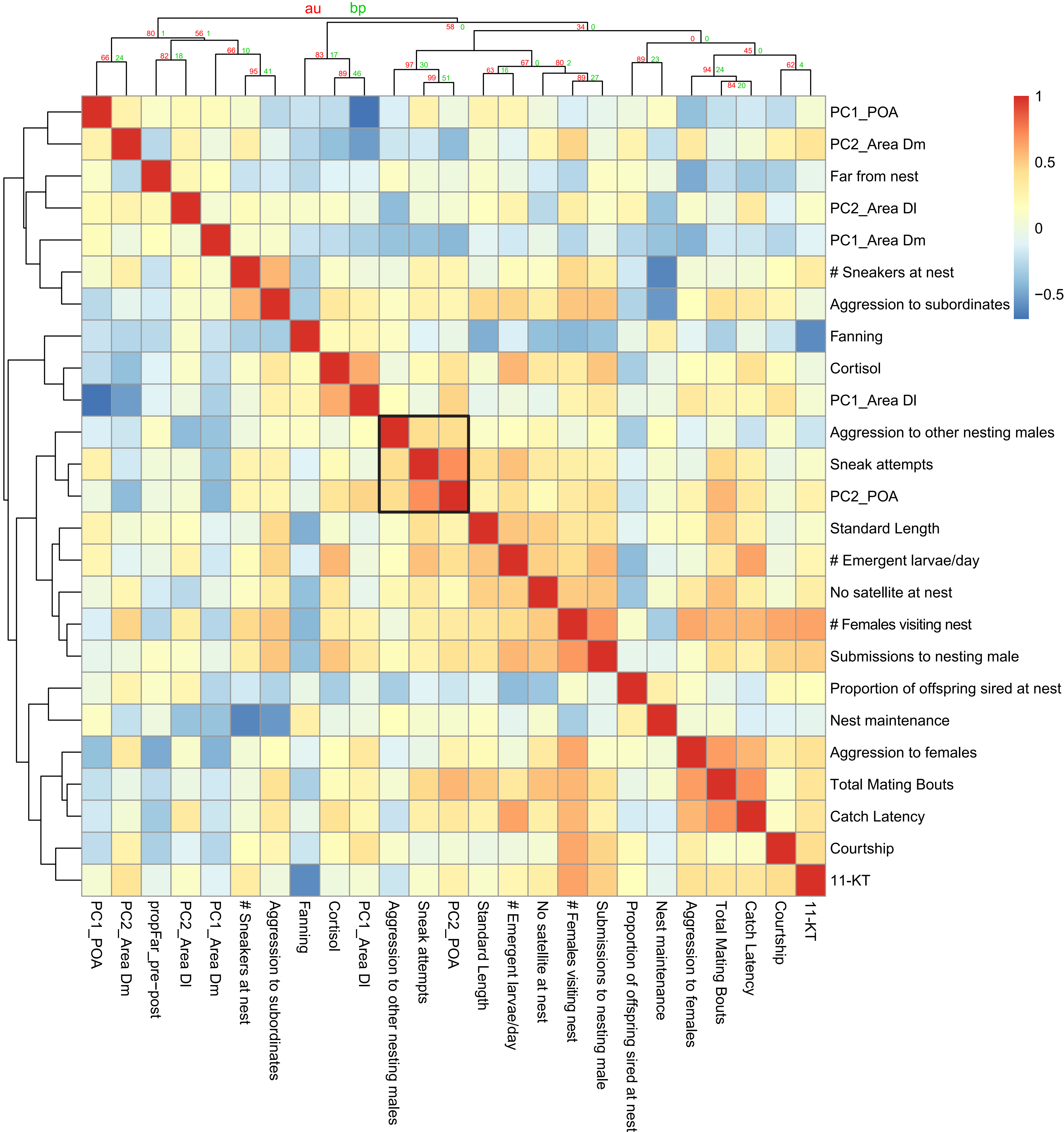
